# Stealth plasmids: rapid evolution of deleted plasmids can displace antibiotic resistance plasmids under selection for horizontal transmission

**DOI:** 10.1101/2025.03.30.646151

**Authors:** Andrew Matthews, Tatiana Dimitriu

## Abstract

Conjugative plasmids carrying antimicrobial resistance (AMR) genes are critical for the spread of AMR, due to their ability to transmit horizontally between bacterial hosts. We previously observed that during experimental evolution in the presence of abundant susceptible hosts, the AMR plasmid R1 rapidly evolves variants with increased horizontal transmission due to mutations causing increased plasmid copy number. Yet AMR was progressively lost from the evolving populations. Here, we show that AMR loss was associated with evolution of ‘stealth’ plasmids in which the AMR region is spontaneously deleted, making plasmid carriage undetectable by plating on selective antibiotic-containing media. These plasmids transmit both vertically and horizontally more efficiently than the ancestral AMR plasmid, driving AMR extinction in bacterial populations and effectively acting as an intrinsic defence against AMR plasmids. Our results suggest that within-host plasmid evolution could be exploited to limit the spread of AMR in natural populations of bacteria.

## Introduction

In bacteria, conjugative plasmids are mobile genetic elements with the ability to transmit horizontally from a donor to a recipient cell, both within and between species. Plasmids play a central role in the dissemination of antimicrobial resistance (AMR) genes among pathogenic bacteria (Dimitriu, 2022; MacLean & San Millán, 2019): a few major plasmid lineages are responsible for the spread of clinically relevant AMR genes, including carbapenamases and extended-spectrum B-lactamases, among Gram-negative bacteria (Carattoli, 2013). Horizontal transmission via conjugation has specifically been implicated in the dissemination of AMR at several scales. For instance at the global scale, world-wide dissemination of the gene *mcr-1* encoding colistin resistance is due to the transmission of a few promiscuous plasmids between strains (Matamoros et al., 2017); and transfer of an azithromycin resistant plasmid facilitated epidemics across multiple *Shigella* species (Baker et al., 2018). Plasmids can also be responsible for the transmission of AMR genes between strains living in different environments, e.g. farm animals and humans (De Been et al., 2014). At the other end of the scale, plasmid transmission occurs between species of clinical enterobacteria within the gut of hospitalised patients (León-Sampedro et al., 2021). Thus, it is crucial to understand what drives horizontal transmission of conjugative AMR plasmids, and what barriers exist to transmission.

Plasmid conjugation depends on both environmental factors (e.g. temperature or spatial structure) and genetic factors (Dimitriu, 2022). It is primarily controlled by the expression of the plasmid-encoded conjugation machinery and its regulatory network (Frost & Koraimann, 2010), but both donor and recipient genotypes also impact conjugation (Sheppard et al., 2020), as well as their relatedness (Dimitriu et al., 2019). Defence systems present in recipients, although likely to have evolved mostly in response to phage predation, also impact conjugation (Dimitriu et al., 2020), and can in turn impact the distribution of AMR genes in pathogens (Pursey et al., 2021). Finally, plasmids themselves can exclude other plasmids, via surface or entry exclusion (Garcillán-Barcia & de la Cruz, 2008), by competition for replication (incompatibility, (Novick, 1987)) or by encoding defence systems targeting plasmids (Benz et al., 2024).

In a previous study, we evolved R1, a model conjugative plasmid conferring resistance to multiple antibiotics, with or without selection for horizontal transmission, and in the absence of antibiotics (Dimitriu et al., 2021). In the presence of potential recipients providing selection for horizontal transmission, plasmids with increased conjugation rate rapidly evolved (Fig. 1A). In most clones, this was due to mutations within the *copA* gene controlling plasmid replication, associated with an increase in R1 plasmid copy number (PCN). We then showed that increased PCN was responsible for an increase in both plasmid conjugation rate and AMR conferred by the plasmid, due to an increase in gene dosage. Despite this, at the population level, there was progressive extinction of AMR plasmids in the treatments with larger daily influx of recipient cells. We interpreted this as a decline in plasmid-carrying lineages due to plasmid horizontal transmission being too low for plasmids to maintain themselves when facing repeated influx of plasmid-free cells. Here, we revisit this study by analysing evolved populations and clones, and perform additional experiments, showing that instead this behaviour is driven by the evolution and invasion of stealth plasmids carrying deletions of the AMR genes, which effectively act as a barrier against AMR plasmids within bacterial populations.

**Figure 1:**
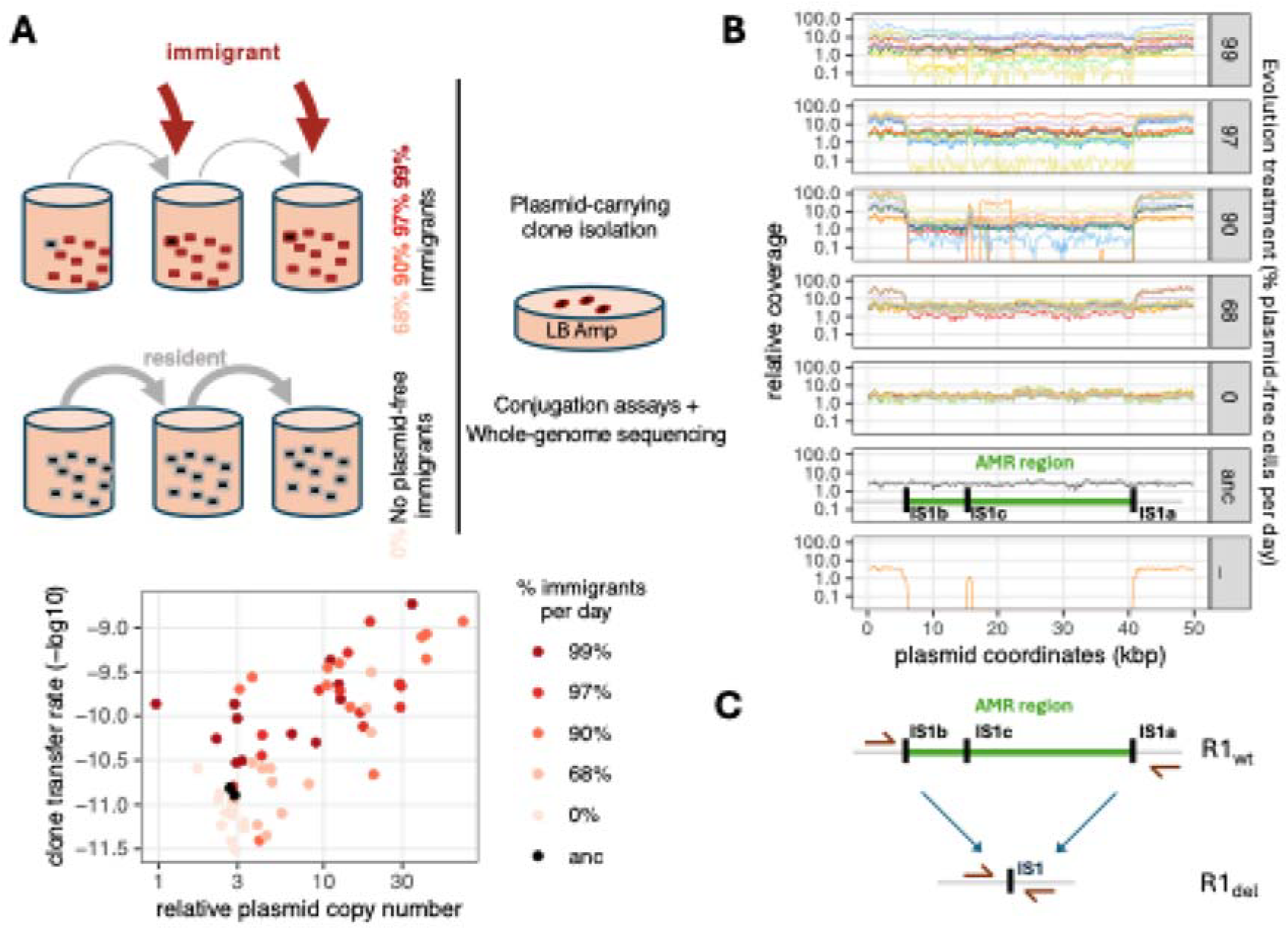
Evolved plasmid-carrying clones with high copy number contain a mix of full-length and deleted plasmids. A: summary of the initial experimental evolution design and phenotypic characterisation of the evolved plasmid-carrying clones. The bottom graph shows transfer rate of evolved clones as a function of relative plasmid copy number measured from sequencing coverage averaged across the plasmid sequence. B: coverage map of evolved clones. Relative coverage of sequencing reads is shown for all clones across R1 sequence length (only the first 50kb are shown). The AMR region and copies of *IS*1 are shown against the ancestral R1 (“anc” treatment) coverage map. C: schematics of the R1_del_ mutations.

## Materials and Methods

### Strains, plasmids and growth conditions

The ancestral strains for our repeat evolution experiment were the same as in our first evolution experiment (Dimitriu et al., 2021): *E. coli* MG1655 (wt) and MG1655 *mutL*::KnR (mut) as initial R1-carrying strains, and wt and mut variants marked with *td-Cherry* as plasmid-free recipients. MG1655 Δ*lac* and a spontaneous rifampicin-resistant mutant of MG1655 (MG1655 Rif^R^) were also used. To test the effect of restriction-modification (RM) systems on transmission, pEcoRI and pEcoRV plasmids, as used in (Dimitriu et al., 2024), were transformed into MG1655 Δ*lac* and maintained using their ampicillin resistance marker. We also used evolved plasmids sequenced from our first evolution experiment, after transfer by conjugation to other host backgrounds. Nomenclature for evolved clones follows (Dimitriu et al., 2021), using bNx, where b = host background (w for wt, m for mut), N = evolution treatment (99, 98, 90, 68 or 0 as % of plasmid-free hosts added at each passage, _ for the no plasmid control treatment) and x = lineage (a to f). ‘Stealth’ plasmid variants with deletion of the full AMR region are indicated as s. Details of strains and plasmids used are shown in Table S1.

Cells were grown in LB medium and clones bearing antibiotic-resistant plasmids were selected with ampicillin 100mg/L. Rifampicin was used at 100 mg/L for selection of MG1655 Rif^R^. *td-Cherry* colonies were identified by their red fluorescence, and Δ*lac* colonies by plating on LB-agar supplemented with IPTG 1mM and X-Gal 0.2g/L. Evolution and competition experiments were done at 37°C, in liquid cultures with shaking at 180 r.p.m., and with 100-fold daily dilution in 200 µL total volume in 96-well plates.

In the repeat evolution experiment, for the immigration treatment, plasmid-free immigrants (wt and mut *td-Cherry* strains) were grown fresh from glycerol stock and mixed with the evolving resident cultures in 95:1 ratio at each passage.

### Detection and characterisation of plasmid variants

Two approaches were used to detect R1 plasmid variants by colony PCR using DreamTaq PCR mastermix (Thermo Scientific). First, primer pairs specific to the backbone of the plasmid (CopA-F & CopA-R, a product indicates that any variant of R1 is present) or to the *bla* gene located in the middle of the AMR region (bla-F & bla-R, a product indicates that the *bla* gene is present) were used for PCR with cycling parameters: 10 min at 95°C, 35 × (15 s at 95°C, 15 s at 52°C, 10 s at 72°C). Second, the R1del-F & R1del-R primers were used for PCR with cycling parameters: 10 min at 95°C, 30 × (15 s at 95°C, 15 s at 64°C, 20 s at 72°C). On the R1 plasmid, these primers are located more than 36kb apart, yielding no product, but they yield a short product when variants with the AMR region deleted are present. Primer sequences are shown in Table S2.

For Sanger sequencing (Eurofins), PCR products were purified with Qiagen QIA Quick.

### Bioinformatic analyses

We reanalysed Illumina sequencing data on evolved clones from (Dimitriu et al., 2021). Unique read coverage from mapping reads to R1 plasmid sequence, generated with breseq as described in (Dimitriu et al., 2021), was averaged over R1 sequence coordinates 1-5570 and 41387-99378 to obtain coverage of R1 backbone; and over R1 sequence coordinates 6200-15335 and 15950-40664 to obtain coverage of R1 AMR region.

### Obtaining bacterial clones carrying stealth plasmids alone

Overnight cultures of evolved clones carrying both full-length and deleted plasmids (based on Illumina sequencing read depth from (Dimitriu et al., 2021), Fig. S1) were mixed with the target recipient strain in a 1:25 donor to recipient ratio during 3h in 500 µL LB broth, then plated without antibiotic selection for AMR plasmid carriage. Recipient background was identified depending on recipient phenotype (red color, white color in presence of IPTG + XGal and rifampicin resistance respectively for *td-cherry*, Δ*lac* and Rif^R^ strains); and colonies containing only deleted plasmids were screened for plasmid presence by PCR (presence of a product with R1_CopA primer pair, absence of a product with R1_bla primer pair) and by checking for ampicillin sensitivity.

### Plasmid invasion and competition experiments

Competitions between R1 plasmid variants were run in 96-well plates. Initial plasmid-carrying clones were chromosomally marked with different colorimetric markers (Δ*lac* or *td-Cherry*, as indicated) to allow detection of transfer to a different strain than the initial donor.

For competitions between variants with vertical transmission, we first conjugated a variant with AMR region deleted into MG1655 Rif^R^ and screened for MG1655 Rif^R^ colonies containing the deleted variant and not containing the full-length (ampicillin-resistant) plasmid. Next, the full-length R1 plasmid (ancestor) was conjugated into this new recipient using ampicillin + rifampicin to select for transconjugants, selecting 4 independent colonies. This was done with evolved clones w_e and w90d, the deleted plasmids s_w_e_ and s_w90d_ being respectively low and high copy number plasmids. Clones were first inoculated into LB containing 50 mg/L kanamycin (Kn) overnight, then cultures were diluted 100-fold every 24 hours into LB without any antibiotic and density of R1-carrying cells was measured using ampicillin resistance as a marker.

For competitions between variants in conditions favouring horizontal transmission, populations were seeded with 1% (vol) overnight cultures of MG1655 containing either R1 or R1_finO_ plasmid (m32e_t_12_, an evolved variant with a *finO* mutation only, see Table S1), 1% overnight cultures of MG Δ*lac* containing either no plasmid or one of s_w_e_, s_w90c_, s_w90d_, s_m68c_ or s_m97f_, and 98% overnight cultures of MG *td-Cherry*. Cultures were then diluted 100-fold every 24 hours into LB without any antibiotic and density of R1-carrying or R1_finO_-carrying cells was measured using ampicillin resistance as a marker. To evaluate the spread of stealth plasmids at 3 days, colony PCR was performed on colonies grown on LB-agar without antibiotics, using the R1del primer pair. Finally, to compare the effect of deleted plasmids and restriction-modification systems as barriers to AMR plasmid invasion, populations were seeded with 1% (vol) overnight cultures of MG *td-Cherry* containing either R1 or R1_finO_ plasmid, and 99% overnight cultures of MG Δ*lac* containing either no plasmid or one of s_w_e_, s_w90d_ evolved plasmids or plasmids carrying EcoRI or EcoRV restriction-modification systems.

### Effect of plasmids on host growth rate

Strains MG1655 Rif^R^ and MGΔ*lac* alone or carrying a single plasmid genotype were grown overnight from glycerol stock without antibiotic selection, then diluted 10,000-fold into 200 µL LB in 96-well microplates and covered with 50 µL mineral oil. Optical density at 600 nm was measured at 5 minute intervals in a Multiskan SkyHigh plate reader at 37 °C for 24h with shaking on. Maximal growth rate was computed using the R package growthrates (Petzoldt, 2018) between 3h and 7h post-inoculation, with the h parameter set at 8. Each plasmid-carrying clone was run along its plasmid-free control, with 14 technical replicates per growth plate and in two different plates (n=28 total).

Data and statistical analysis used R version 4.3.2 (R Development Core Team, 2008).

## Results

### Existence of deleted plasmids within AMR clones

Previously, we studied R1 plasmid evolution in conditions that favour horizontal transmission ((Dimitriu et al., 2021), Fig. 1A). We sequenced 60 plasmid-carrying clones isolated by plating evolved populations in the presence of ampicillin, and identified multiple plasmid variants carrying point mutations in the *copA* gene (summarized as *copA\** mutations), associated to an increased plasmid copy number (PCN), which caused increased horizontal transmission via gene dosage effects (Dimitriu et al., 2021). Increased PCN of *copA\** plasmids was apparent as increased coverage of mapped reads when mapping reads from Illumina sequencing to the ancestral plasmid sequence. However, we also noticed that the depth of coverage of sequencing reads is not uniformly high along the length of R1 plasmid sequence (Fig. 1B, Fig. S1): instead, for many plasmid variants there is a drop in sequencing depth approximately between coordinates 5700 and 41300. Depending on the clones, read coverage drops to values which are either higher than the ancestral plasmid, equal to the ancestral plasmid (2-3 copies per chromosome), or lower than the ancestral plasmid or even than the chromosome (Fig. 1B). We hypothesized that this variation in sequencing depth is due to a deletion present within R1 sequence, in some of the plasmid copies. Strikingly, the low coverage region includes all antibiotic resistance genes present on R1, including the *bla* gene conferring ampicillin resistance. As we initially isolated R1-carrying clones using ampicillin, the clones we sequenced could not have fully lost the resistance determinant, and the result is a mixture of R1 sequence variants co-existing within each bacterial clone (likely facilitated by the evolution of high copy number).

To identify and characterise these deleted plasmids, we took advantage of the plasmid-free control populations from our evolution experiment, for which we had sequenced clones without any antibiotic selection step. In one clone, w_e, sequencing revealed the existence of reads mapping to R1, which must have originated from contamination, likely from the nearby evolving plasmid-carrying populations. Here, no reads mapped within the resistance determinant, suggesting a full deletion (Fig. 1B bottom). We designed PCR primers around this region, which yielded a product just above 1kb long in this clone, confirming a large deletion exists within the variant plasmid (Fig. S2). From now on, we call deleted variants R1_del_. Interestingly, while there is no PCR product in the absence of R1 plasmid, clones carrying the ancestral R1 plasmid also yield a faint band of the same size (Fig. S2), suggesting that the variant occurs spontaneously at low frequencies in R1 colonies, although some PCR artefact is also possible. We next sequenced the PCR product using Sanger technology to characterize the deletion in more detail. The approximately 35.6kb long deleted region is bounded in the ancestral plasmid by two identical insertion sequences (IS) IS1 copies, IS1b and IS1a, in direct orientation. Sanger sequencing revealed that the R1_del_ variant sw_e experienced a deletion bounded exactly by the two external IS sequences IS1b and IS1a. The deleted plasmid retains one unique IS copy, which suggests it evolved by homologous recombination between the two initial direct repeats (Fig. 1C).

Coverage data shows that R1_del_ variants are also present in many clones selected using ampicillin, coexisting with the full-length plasmid, and leading to uneven read depth along the plasmid sequence (Fig. 1B). To quantify this, we compared short-read coverage between the AMR region and the rest of the plasmid, or plasmid backbone (Fig. S3A). While a few evolved clones have similar coverage between the AMR region and the plasmid backbone, the majority of high PCN clones have higher coverage for the backbone, with many of them showing no increased coverage for the AMR region compared to the ancestor. This suggests that the majority of these clones contain one version of the plasmid similar to the ancestor R1_wt_, together with a high copy number, R1_del_ plasmid. Accordingly, the large majority of clones with the *copA\** allele (high PCN variants) have a relative coverage of the AMR region around 10% of the coverage of the plasmid backbone (Fig. S3B), meaning that in these clones only deleted plasmids have high PCN, and the AMR region is part of a plasmid molecule with ancestral PCN.

### AMR loss is associated with the spread of deletion plasmids

We previously followed plasmid population dynamics using the ampicillin resistance phenotype conferred by the *bla* gene within the AMR region in the full-length plasmid and concluded that plasmids went extinct. Identification of deleted variants suggests that instead, AMR plasmids went extinct but R1_del_ plasmids might have survived for longer and escaped detection in antibiotic-sensitive clones. Consistent with AMR plasmids being subject to evolutionary rather than direct ecological dynamics is the observation that AMR population density began declining only after 12 to 15 days. If ecological dynamics alone drove this decline (due to insufficient horizontal transmission), we would expect a steady trend beginning much earlier. To test the robustness of this delayed AMR decline, we first repeated a subset of our previous experiment (adding either no recipient or plasmid-free recipients representing 95% of the total bacterial population every day) for 32 days. We found that the population dynamics in our repeat evolution experiment was very similar to the first (Fig. 2A) with AMR cell density beginning to decline below its initial levels after approximately 10 days in the presence of plasmid-free recipients. Interestingly, this time we observed a later decline in AMR cell density even in the absence of plasmid-free recipients.

**Fig. 2:**
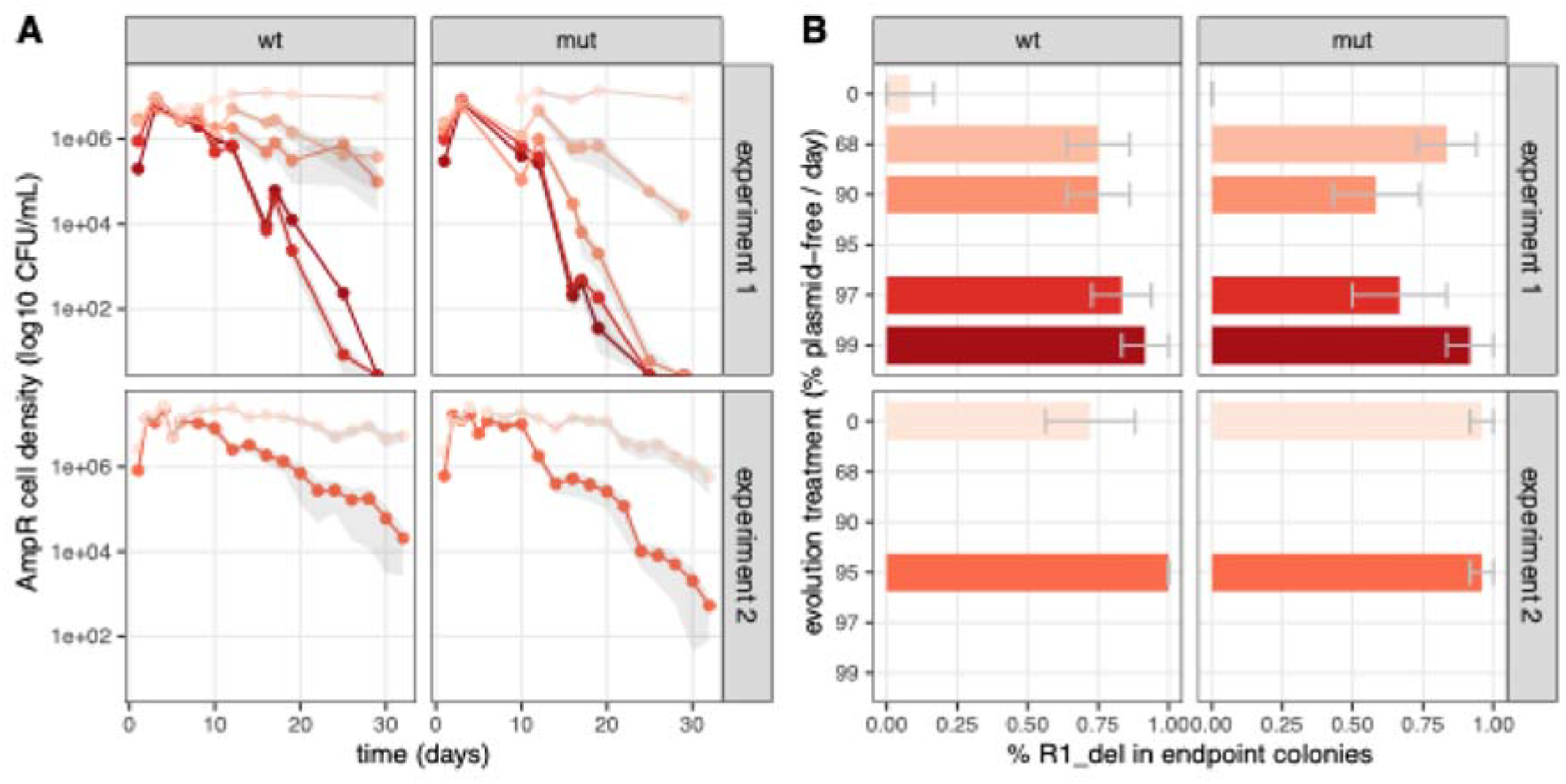
AMR declines repeatedly during evolution in the presence of plasmid-free host bacteria, but stealth plasmids are maintained at high density. A shows the dynamics of ampicillin resistant cell density in two independent experiments for two host backgrounds (wt and mut). B shows endpoint frequencies of R1_del_ carriage. Colour and y-axis in B indicate evolution treatment. Each treatment x host background combination was run in 6 independent evolution replicates; mean values per conditions are shown in colour and mean ± s.e.m. is shown in gray, as a shaded area in A and as an error bar in B.

In light of our results on individual clones, we revisit our analysis of plasmid population dynamics data. We hypothesize that plasmids might have persisted for the full length of the experiment but escaped detection using antibiotic resistance as a marker, due to deletion of antibiotic resistance determinants. We used PCR with the R1del primer pair to discriminate between full-length and deleted plasmid carriage (see Fig. S2), and screened colonies obtained from reviving frozen endpoint populations on LB-agar in the absence of antibiotic selection. Strikingly, we found that R1_del_ variants are present in most colonies tested for all populations evolved with regular influx of plasmid-free cells. In the second experiment, in which we observed a late decline in AMR cell density even in treatments without regular influx of plasmid-free cells, endpoint populations for these treatments also contained a large proportion of R1_del_ variants. Thus, the observed decline in AMR cell density is not due to plasmids being lost from host populations; instead R1_del_ variant ‘stealth plasmids’ evolve and are efficiently transmitted within populations. This suggests that the decline in AMR full-length plasmids might be itself due to displacement by R1_del_ variants.

### Stealth plasmids transmit more efficiently than AMR plasmids

To understand why stealth plasmids displace full-length AMR plasmids at the population level, we next studied their vertical and horizontal transmission.

We analysed vertical transmission of the ancestral R1 plasmid (using its ampicillin resistance phenotype to follow R1-carrying cell density) in clones where it co-exists with a R1_del_ variant at the start of the experiment. We focused on two deleted variants varying in their PCN: s_w_e_, which has an ancestral, low PCN, and s_w90d_, which evolved high PCN. Over 8 days, R1 is maintained in the population with no detectable loss when competing with a R1_del_ variant with low PCN. In contrast, it declines rapidly when competing with a R1_del_ variant with high PCN (Fig. 3B). Thus, the full-length ancestral R1 does not appear to experience a vertical transmission disadvantage against deleted variants purely because of plasmid size, but high-copy number variants (which represent the majority of deleted variants present in sequenced clones, see Fig. 1C) rapidly displace it.

**Fig. 3:**
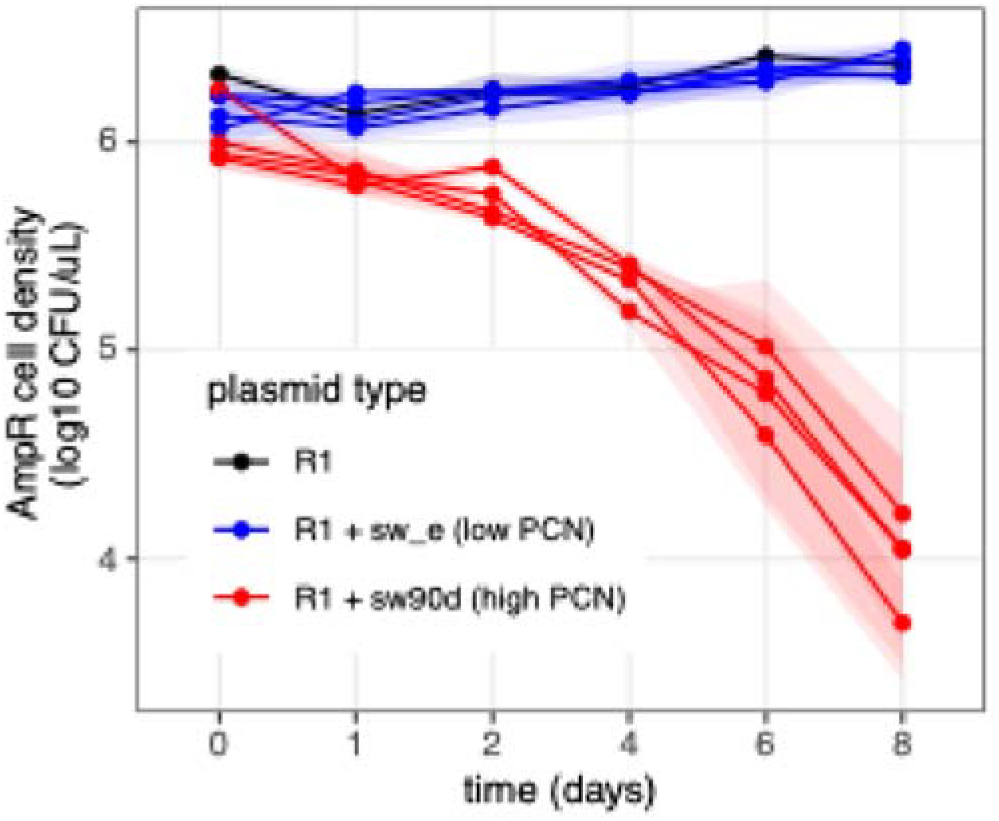
The outcome of competition at vertical transmission depends on plasmid copy number. R1-carrying cell density of populations seeded with clones containing R1 alone or co-existing with stealth variants is shown as a function of time. Each line represents an independent founding clone, and shaded areas show standard error from 4 replicate experiments; colour indicates initial plasmid content.

Next, to evaluate horizontal transmission and its contribution to plasmid spread, we performed an experiment in which stealth plasmids were initially present in a different clone than R1 plasmid, both at low density in the presence of abundant potential recipients. In these conditions, overall plasmid dynamics will be driven to a large extend by horizontal transmission. Here we used evolved stealth plasmids that differ in their PCN *copA\** mutations, including one which is also a *finO* derepressed mutant, as well as AMR variants R1 and R1_*finO*_, as we expect this to impact horizontal transmission.

In control populations without stealth plasmid, R1 invades the cell population in 3 to 4 days, whereas R1_*finO*_ has already invaded after 1 day (Fig. 4A top, black). In the presence of all stealth plasmids, R1 spread is stopped or strongly limited at all time points (Fig. 4A top, other colours). This includes the s_w_e_ variant which has an ancestral PCN and no *finO* mutation, thus no obvious mutation providing a transmission benefit against R1 apart from the AMR region deletion. In contrast, the presence of most stealth plasmids has little effect on R1_*finO*_ spread in the population at early timepoints, with only s_m68c_ variant limiting AMR densities (Fig. 4A bottom, dark blue). s_m68c_ carries a *finO* mutation similarly to R1_*finO*_, suggesting that only a stealth plasmid with transfer derepression ‘matching’ AMR plasmid transfer derepression can significantly impact early spread. Indeed, focusing on the dynamics of AMR in the recipient (initially plasmid-free) cell population shows the same pattern as for all AMR cells (Fig. 4B), demonstrating that the observed dynamics is due to horizontal transmission to the larger recipient population. Interestingly, stealth plasmid inoculation still influences R1_*finO*_ spread but at later time points, with a decrease in R1_*finO*_ population density overall and in transconjugants, suggesting this is driven by the large cost R1_*finO*_ imposes on its host rather than by differences in horizontal transmission. Moreover, screening colonies after 3 days for the presence of stealth plasmids shows that in competition with the slower R1, all stealth plasmids, have already invaded recipients, whereas in competition with R1_*finO*_, horizontal transmission of stealth plasmids is more limited (Fig. 4C).

**Fig 4.**
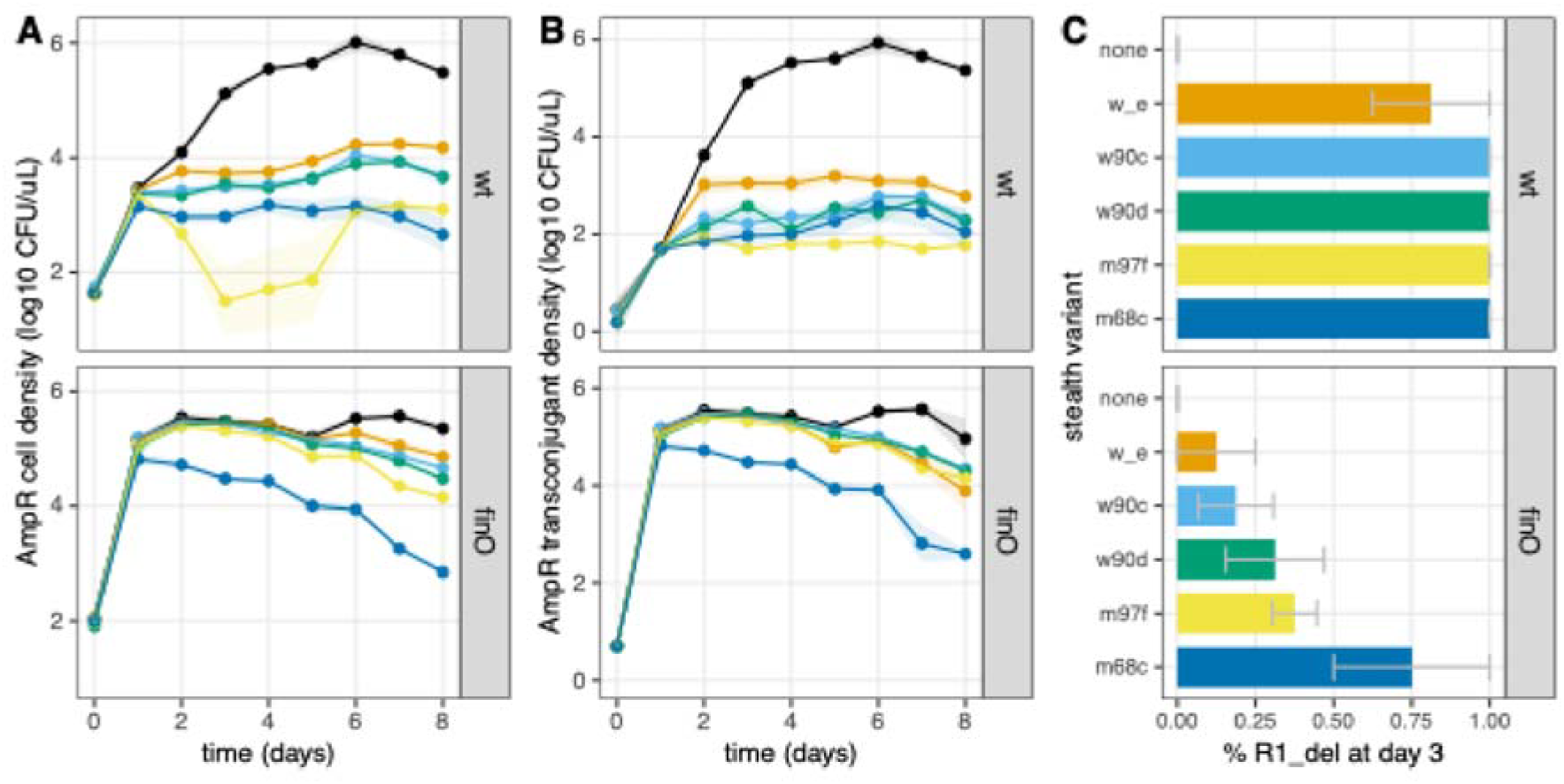
Stealth plasmids invade horizontally faster than ancestral plasmids. Populations were seeded with 1% AMR plasmid-carrying cells, 1% stealth plasmid-carrying cells (variant indicated by colour and detailed in C), and 98% plasmid-free cells. Total AMR cell density is shown in A, and density of AMR transconjugants is shown in B, with lines and shaded areas indicating respectively the average and standard error from 4 replicate experiments. C shows the proportion of clones carrying stealth plasmids at day 3 for each stealth treatment (4 colonies tested for each of 4 replicate experiments, error bars show standard error across replicate experiments).

Thus, these experiments show that all stealth plasmids transmit faster than the full-length R1 AMR plasmid and readily prevent its spread to recipient cells.

### Cost to the host is determined mostly by transfer gene expression

To measure how plasmid variants affect the growth of their hosts, we measured the exponential growth rate of two strains (MG Rif^R^ and MG Δ*lac*, both used in the previous experiments) alone or carrying R1 or one of R1’s evolved variants (Fig. 5). MG Rif^R^ grew overall more slowly than MG Δ*lac*, and plasmid type had a significant effect on growth rate, which also depended on the host (growth rate ∼ strain * plasmid, strain effect F_1,324_=330, p<2 10^−16^, plasmid effect F_7,324_=76.9, p<2 10^−16^, interaction effect F_3,324_=0.0015).

**Fig. 5:**
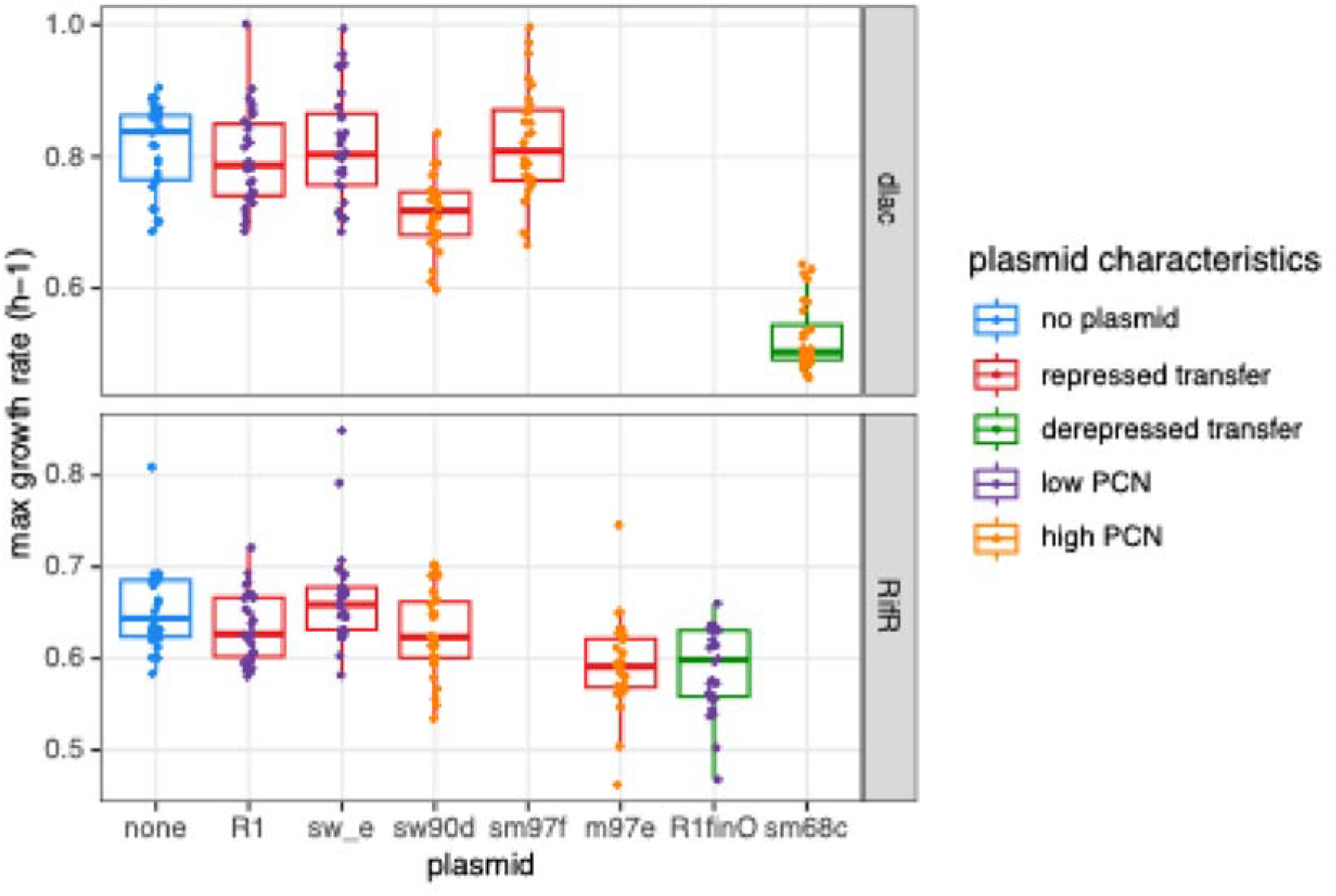
Copy number and mutations in transfer regulation impact stealth plasmid cost. Exponential growth rate is shown for plasmid-free and plasmid-carrying cells in two strain backgrounds, MG Δ*lac* (top) and MG Rif^R^ (bottom). The centre line of the boxplots shows the median, boxes show the first and third quartile, and whiskers represent 1.5 times the interquartile range; individual data points are shown as dots (n=14). Boxplot colour indicates if transfer is repressed (based on plasmid *finO* genotype), and dot colour indicates PCN (based on plasmid *copA* genotype).

Neither the ancestral R1 plasmid or the low PCN s_w_e_ had a detectable effect on growth rate (see Table S3 for detailed statistics). The low PCN stealth variants s_w90d_ and s_m97f_ had a variable effect on growth, with no effect of s_m97f_ in MGΔ*lac* and no effect of s_w90d_ in MG Rif^R^, but a significant effect of s_w90d_ in MG Δ*lac* (TukeyHSD test, -0.10±0.05 h^-1^ vs plasmid-free clone, p=6.8 10^−6^, -0.082±0.05 h^-1^ vs R1-carrying clone, p=0.0003). We additionally tested in MG Rif^R^ the effect of m97e plasmid, one of the three only evolved R1 variants that had uniformly high PCN (with no AMR region deletion). m97e’s cost on growth was significant (−0.061±0.036 h^-1^ vs plasmid-free clone, p=3.7 10^−5^), and higher than the cost of a stealth high copy variant in the same strain (−0.037±0.036 vs s_w90d_, p=0.04), showing that deletion of the AMR region limits the cost of high PCN plasmids. Finally, we also measured the cost of the R1_finO_ variant which displays derepressed transfer, as well as the cost of s_m68c_, a stealth variant which has both high PCN and a similar *finO* mutation leading to derepressed transfer. We found that R1_finO_ also imposed a significant cost to its host MG Rif^R^ (−0.065±0.036 h^-1^ vs plasmid-free clone, p=9.3 10^−6^, -0.046±0.036 h^-1^ vs R1-carrying clone, p=0.005); and s_m68c_ imposed a much higher cost again to its host MG Δ*lac* (−0.29±0.05 h^-1^ vs plasmid-free clone, p=1.1 10^−13^, -0.27±0.05 h^-1^ vs R1-carrying clone, p=1.1 10^−13^). This cost is also significantly higher in MG Δ*lac* than the cost of s_w90d_ (−0.19±0.05, p=1.6 10^−13^), which has high PCN but repressed transfer operon expression. s_m68c_ displaying the highest cost is consistent with transfer gene expression placing a high burden on the host, as high PCN will multiply the effect of increased gene expression from *finO* mutations. Thus, overall plasmid carriage cost appears to be determined first by the level of transfer gene expression, followed by PCN, leading to the AMR region deletion relieving some fitness cost specifically in high PCN plasmids.

### Stealth plasmids as an efficient barrier to AMR plasmids

The effect of stealth plasmids limiting AMR plasmid transmission, observed here, implies that acquisition of stealth plasmids by susceptible cells makes these cells resistant to further plasmid entry. This effect is expected due to entry exclusion functions commonly expressed by plasmids (Garcillán-Barcia & de la Cruz, 2008) including R1 (Cox & Schildbach, 2017). To focus on this effect, we now directly test how stealth plasmid presence in recipient cells affects R1 conjugation and compare their effect with that of canonical defence systems, restriction-modification (RM) systems (Fig. 6). We include 3 stealth plasmids differing in relevant transmission phenotypes: s_w_e_ (low PCN), s_w90d_ (high PCN) and s_m68c_ (high PCN, derepressed transfer gene expression); and two Type II RM systems that were among the most effective against plasmid conjugation in our previous study (Dimitriu et al., 2024).

**Fig. 6:**
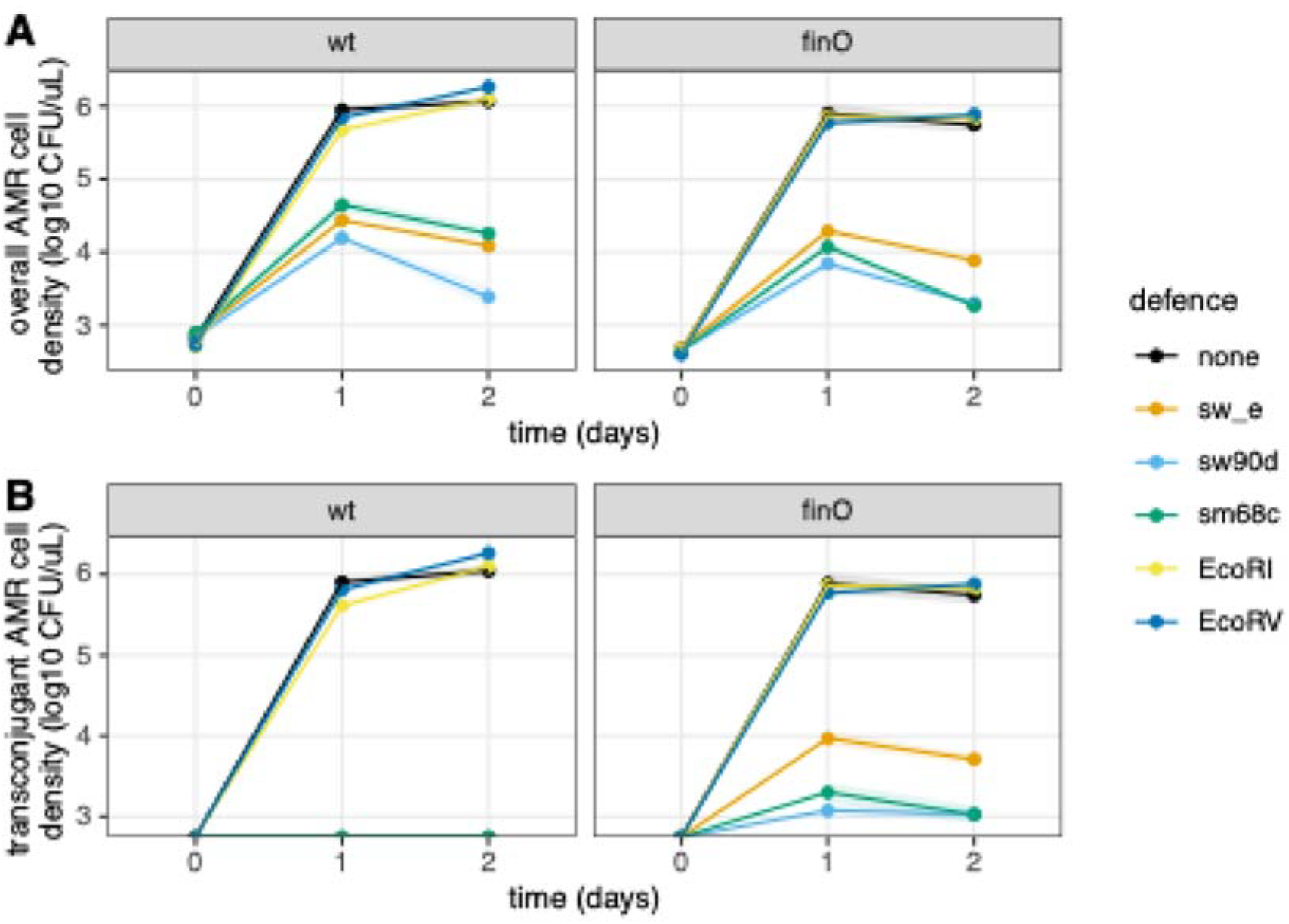
Stealth plasmids are a more efficient barrier to AMR plasmids than RM defence systems. Populations were seeded with 1% AMR plasmid-carrying cells (R1 on the left, R1_*finO*_ on the right) and 99% AMR-free cells, either with no defence system or with one of s_w_e_, s_w90d_ or s_m68c_ plasmids or EcoRI or EcoRV RM systems. Total AMR cell density (A) and AMR cell density in initially AMR-free cells (B) are shown as a function of time, with lines and shaded areas indicating respectively the average and standard error from 4 replicate experiments.

Strikingly, we find that all 3 stealth plasmids strongly limit the spread of both R1 and R1_*finO*_ in the mixed population, whereas the two RM systems have no significant effect on plasmid spread (Fig. 6A). Moreover, the smaller increase in cell density in the presence of stealth plasmids is due exclusively to donor cell growth in the case of R1, for which no transconjugants were detected in any of the stealth plasmid-containing recipients; and even the spread of the more transmissible R1_*finO*_ is due primarily to horizontal transmission only in the presence of the low PCN s_w_e_ plasmid, but not in the presence of high PCN s_w90d_ or s_m68c_ (Fig. 6B). This suggests that high PCN of the stealth plasmids favours their exclusion of AMR plasmids. Overall, we conclude that stealth plasmids are more efficient as a barrier to the AMR plasmids they are evolved from, compared to RM defence systems.

## Discussion

Previously, we analysed the ecological and evolutionary dynamics of R1 plasmid exposed to regular immigration of plasmid-free recipients. Using antibiotic resistance as a phenotypic marker for R1 carriage, we observed evolution of high plasmid copy number, associated with increased conjugation rate and antibiotic resistance. Yet, R1 population size still declined over time in the face of cell immigration. Here we show that this decline in resistance is due to rapid evolution of stealth plasmids with deletions of the AMR region (undetectable by standard assays using resistance to antibiotics to detect plasmid carriage), which displace ancestral resistance plasmids. Thus, at the population level evolution ultimately leads to AMR extinction despite a transient increase in AMR associated to high PCN.

To understand this outcome, we studied the dynamics of R1-carrying cells when exposed to stealth plasmid variants. At vertical transmission, high PCN stealth variants rapidly displace R1. This is expected as plasmids are competing not only for replication itself but also for partitioning to daughter cells, and early experiments showed that high PCN mutants of R1 displaced R100, another IncFII plasmid, faster than R1 did (Nordström et al., 1972). In contrast, R1 seems to be unaffected by the low PCN s_w_e_ variant. This demonstrates a direct effect of high PCN on vertical transmission; however, our assay has low sensitivity and low PCN plasmids might still have some significant, if lower, role in R1 displacement. During horizontal transmission, all the stealth variants we tested - including the low PCN s_w_e_ - spread faster than the ancestor AMR plasmid. A smaller plasmid size might directly be responsible for increased horizontal transmission, possibly because it allows conjugative replication itself to complete faster. Still, we must note that s_w_e_ also carries another mutation, a G deletion in a polyG tract within the leading region (Table S1). This mutation was frequent across evolved clones (Dimitriu et al., 2021) and we cannot exclude that it contributes to s_w_e_’s horizontal transmission benefit. High PCN stealth plasmids still spread faster than s_w_e_, in accordance with our previous results showing that high PCN leads to higher conjugation rates (Dimitriu et al., 2021). Finally, plasmid population dynamics will also be affected by plasmid cost to the host. Growth rate assays show that stealth plasmids have a limited cost, mostly dependent on other mutations present in the evolved plasmids – with mutations leading to derepression of transfer being particularly costly, especially when combined with high PCN. Comparing a high-copy stealth variant and a high-copy AMR variant showed a lower cost of the stealth variant suggesting that at least in high-copy plasmids, AMR gene deletion will bring fitness benefits to the host (and in turn to plasmids themselves via increased vertical transmission). We did not observe any cost for the ancestral R1 plasmid, preventing the detection of any effect of the deletion on plasmid cost in low-copy variants. Yet, deletions of AMR genes can improve plasmid cost in other low-copy plasmids, in particular in the case of highly-expressed □-lactamases (DelaFuente et al., 2022; Humphrey et al., 2012; Rajer & Sandegren, 2022). Previous studies have shown that both within- and between-host competition contribute to plasmid fitness (Hülter et al., 2020; Rossine et al., 2025). Here, we show that in conjugative plasmids, variants can compete for vertical as well as horizontal transmission, in addition to selection happening both within and between cells.

In our evolution experiment, we observed deletions in almost all clones with high PCN, suggesting R1 has a high propensity to experience deletions. Indeed, previous evolution experiments with R1 observed deletions in a similar immigration design (Smith, 2011) or at longer timescales in the absence of immigration (Dahlberg & Chao, 2003). Early molecular studies also showed that R1 and other IncF plasmids can spontaneously dissociate into elements corresponding to its backbone and its resistance region, due to recombination between the IS1 sequences (Chandler et al., 1977; Silver & Falkow, 1970). More generally, ISs are generally abundant on plasmids, and specifically enriched in conjugative plasmids encoding AMR genes, with the regions surrounding AMR genes being particularly dense in ISs (Che et al., 2021). Deletions of AMR genes due to recombination between ISs might thus be a general phenomenon across conjugative plasmids, and have been observed experimentally e.g. (Lee & Marx, 2012; Zhang et al., 2022). Chromosomal rearrangements leading to deletions can also arise from IS recombination (Raeside et al., 2014). However, plasmids might be particularly prone to deletions, due precisely to their richness in large DNA repeats and presence in cells at high copy number (Cury et al., 2018). In our experiments, high PCN likely evolved first - being selected due to its benefits for horizontal transmission - which then promoted recombination and evolution of stealth plasmids. The emergence of deleted variants within cells that still contain full-length variants then likely leads to complex eco-evolutionary dynamics with extended periods of co-existence, as observed recently in (Rossine et al., 2025). It remains to be determined how within-cell and between-cell selection interact, and how long high PCN variants remain stable.

Overall, stealth plasmids invade populations more efficiently than their AMR ancestor and when present in a recipient cell, they act as an effective barrier to AMR plasmid conjugation. This barrier effect likely arises mostly from entry exclusion by plasmids present in the recipient cell (Garcillán-Barcia & de la Cruz, 2008). In the F plasmid, which transfer machinery is similar to R1’s, entry exclusion decreases transfer 100-to 300-fold (Achtman et al., 1977). This could explain how the initial faster spread of stealth variants via horizontal transmission translates into lasting limitation of AMR spread. By contrast, plasmids rapidly escape restriction by two RM systems. This is consistent with our previous results, in which median plasmid restriction during short-term conjugation assays across plasmids and RM systems in *E. coli* was only 14-fold (Dimitriu et al., 2024). Plasmids have evolved multiple anti-defence mechanisms (Tesson et al., 2024) as well as defence avoidance strategies (Shaw et al., 2023). Avoiding competition arising from core replication mechanisms or from entry exclusion encoded by the plasmid immediate ancestor is likely harder to achieve. Moreover, host defences are likely under little selection pressure to evolve to target plasmids (much less costly to host cells than phages), whereas plasmids have evolved to compete with each other for host resources (Rocha & Bikard, 2022). AMR and stealth plasmids are effectively competing for the host niche, translating into exclusion of AMR plasmids by stealth ones, similar at a lower selection level to the replacement of AMR strains by competing non-AMR ones in fecal transplants (Woodworth et al., 2023).

Our results suggest that plasmid competition might serve to limit the spread of mobile AMR, whether in the context of natural populations, or through engineering approaches. AMR plasmids can cohabit naturally with non-AMR, highly related plasmids within populations (Lipworth et al., 2024), providing opportunities for competitive exclusion dynamics. For instance, in a genomic study of bloodstream isolates, an antibiotic-susceptible lineage was found to contain a non-AMR IncF plasmid related to the AMR plasmids carried by other, resistant isolates, suggesting that the susceptible plasmid excluded AMR plasmids (Stephens et al., 2017). In a CRISPR-based engineering approach, supplying a mobilizable plasmid incompatible with the target plasmid also helped prevent the spread of AMR, especially in conditions of horizontal transmission (Wongpayak et al., 2021). Stealth plasmids evolving from AMR plasmids would have the added benefit of being antibiotic-sensitive, so do not risk disseminating other marker genes into the environment. Finally, at longer timescales evolution of deleted plasmids and their displacement of competing plasmids carrying various accessory genes could contribute to the pattern of accessory gene depletion recently observed in plasmids compared to chromosomes (MacLean et al., 2025) and shape the broader mobile gene pool.

## Data availability

All data underlying this work are available in the article and in its online supplementary material.

