## Supplementary Tables and Figures for "Stealth plasmids: rapid evolution of deleted plasmids can displace antibiotic resistance plasmids under selection for horizontal transmission"

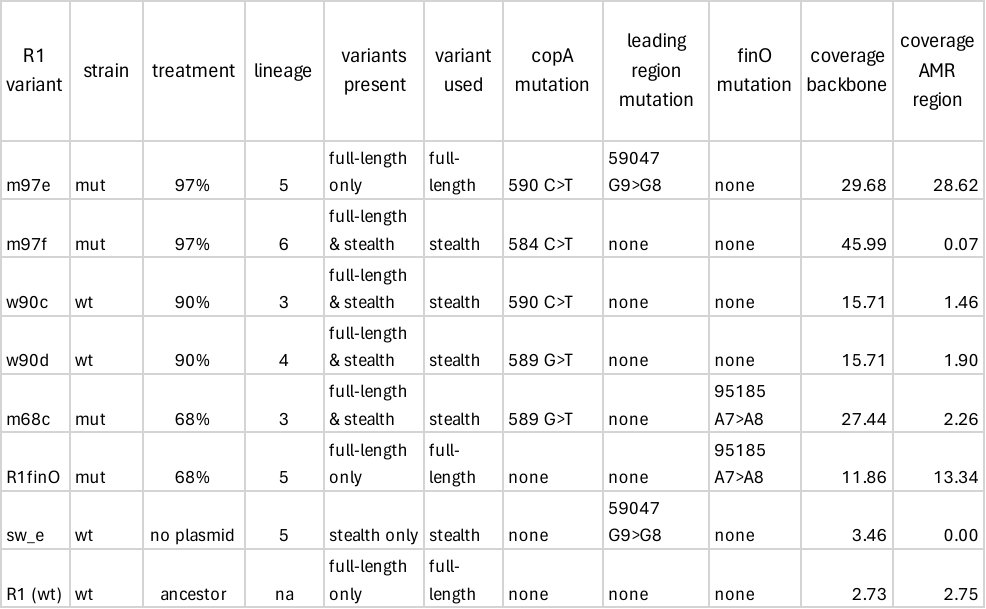

**Table S1: R1 variant plasmids used in this study.** Mutations present in R1 variants are indicated.

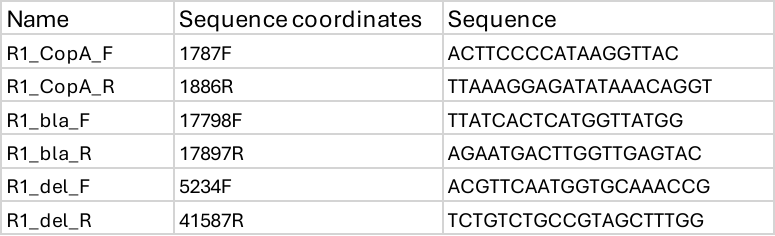

**Table S2: primers used in this study**

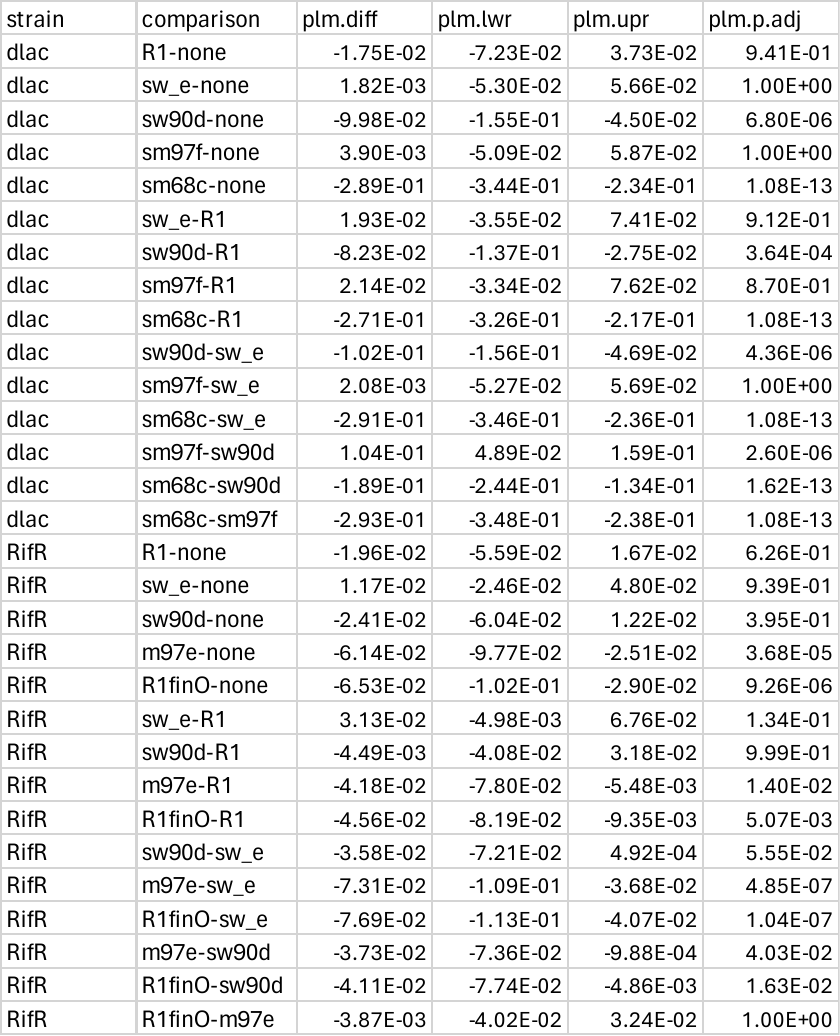

**Table S3: statistical analysis of growth rate data.** Results of TukeyHSD tests performed separately for each host strain are shown for all plasmid type comparisons.

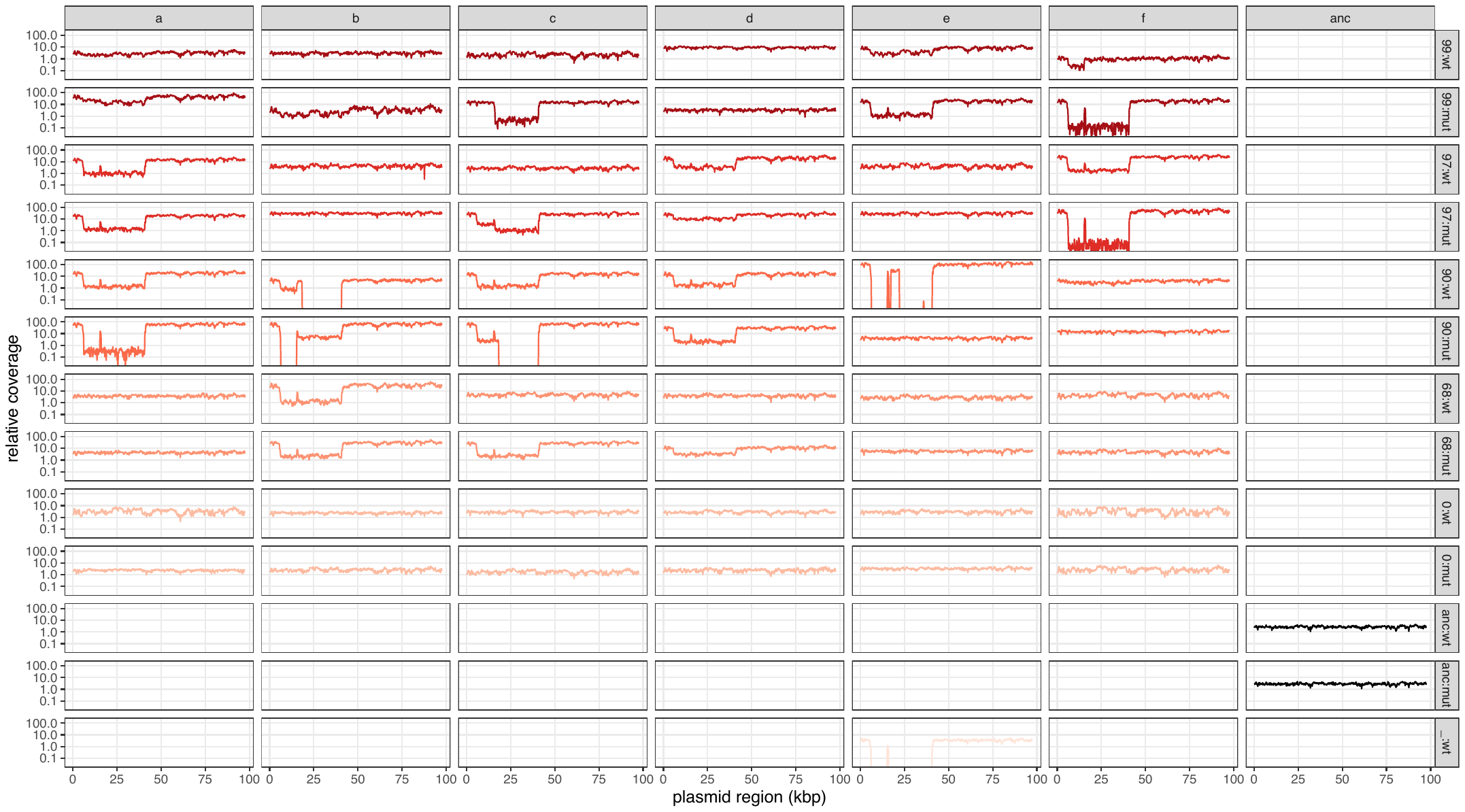

**Fig. S1: Detail of Illumina sequencing read coverage across R1 plasmid sequence**. Relative coverage of sequencing reads is shown for all clones across R1_wt_ sequence map. Relative coverage was measured as the sum of coverage of both unique and repeat reads, divided by the overall average coverage of reads mapped to the chromosome.

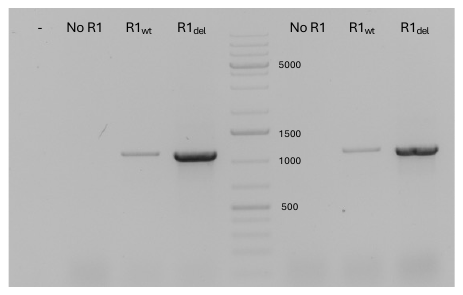

**Fig. S2: Agarose gel electrophoresis of the products of colony PCR using R1_del_ primers*.*** “-“ indicates the negative control without bacteria; other reactions were run with *E. coli* carrying no R1, R1_wt_ or R1_del_ as indicated, in two replicates from independent colonies. Numbers indicate the size of some DNA ladder bands, in bp.

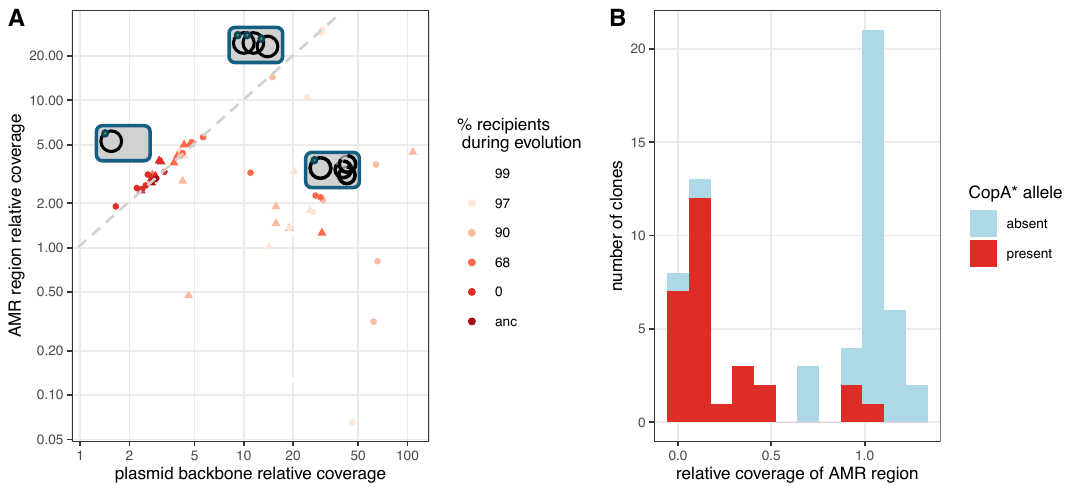

**Fig. S3**: **Characterisation of coverage variation along sequenced evolved R1 clones.** In A, short-read coverage of the AMR region is shown on the y-axis and coverage of the rest of the plasmid is shown on the x-axis; each dot represents a sequenced clone with evolution treatment indicated by color and strain background by dot type (circles = wt, triangles = mut). Interpretation of plasmid content is shown for three regions of the graph, with bold lines showing plasmid regions present, thin lines deleted regions, and the green circle indicating the ampicillin resistance marker. B shows the distribution of relative coverage of the AMR region (compared to the backbone region), with *copA** plasmids in red and ancestral *copA* plasmids in light blue.
